# Mechanical models affecting beetle horn remodeling

**DOI:** 10.1101/2023.01.16.524208

**Authors:** Keisuke Matsuda, Haruhiko Adachi, Hiroki Gotoh, Yasuhiro Inoue, Shigeru Kondo

## Abstract

Clarifying the mechanisms of shape alteration by insect metamorphosis is important for comprehending exoskeletal morphogenesis. The large horn of the Japanese rhinoceros beetle *Trypoxylus dichotomus* is the result of drastic metamorphosis, wherein it appears as a rounded shape via pupation and then undergoes remodeling into an angular adult shape. However, the mechanical mechanisms of this remodeling process remain unknown. In this study, we investigated the remodeling mechanisms of the Japanese rhinoceros beetle horn by developing a physical simulation. We identified three factors contributing to remodeling by biological experiments—ventral adhesion, uneven shrinkage, and volume reduction—which were demonstrated to be crucial to the transformation by a physical simulation. We also corroborated our findings by applying the simulation to the stag beetle’s mandibular remodeling. These results indicate that the physical simulation is applicable to pupal remodeling in other beetles, and the morphogenic mechanisms could explain various exoskeletal shapes.

**Significance statement:** The metamorphosis in insects is a mysterious process. By metamorphosis, insects sometimes change their shape dramatically. The head horn of the Japanese rhinoceros beetle is one of the most famous examples of metamorphosis. In larva-to-pupa molting, the horn appears suddenly, caused by the “furrow formation and unfolding” mechanism. The unfolding process makes the pupal horn rounded. However, pupa-to-adult molting transforms the rounded shape into an angular shape. In this paper, we investigated the mechanisms of the transformation. We extracted factors contributing to it through observations and experiments and developed a physical simulation. It could reproduce the adult shape from the pupal shape and could be a general model for the pupa-adult transformation of beetles.

## Introduction

Understanding the mechanisms of animal morphogenesis is challenging, particularly in exoskeletal animals with great morphological diversity. These animals grow discontinuously through molting, which enables some hemimetabolous and holometabolous insects to undergo dramatic changes in shape, known as metamorphosis (1). Exoskeletal animals are covered with a rigid cuticle layer, whose undulation determines the animals’ shape. Typically, cuticle shaping is achieved as follows: the pre-existing epithelial sheet just beneath the current cuticle (the outermost layer) detaches from the cuticle through apolysis in preparation for molting. Cellular (e.g., cell division) and physical factors (e.g., internal pressure) then alter the epithelial and cuticle layers before and after ecdysis, resulting in a different form of the cuticle layer(1–3). Nonetheless, the control mechanisms for the shape of the layers remain poorly understood.

From a physical standpoint, force distribution determines the epithelial sheet deformation (4–6). The factors controlling force distribution can be divided into cell-intrinsic and cell-extrinsic factors. Cell-intrinsic factors, such as the cytoskeleton and cell–cell junctions, generate short-range force patterns, while cell-extrinsic factors, such as extracellular matrices (ECMs), generate long-range force patterns (4, 7). While the effects of cell-intrinsic factors on morphogenesis have been extensively studied in developmental biology, the effects of cell-extrinsic factors have only recently begun to be investigated. For instance, one study examining the role of cell-extrinsic factors on cell sheet deformation in *Drosophila* reported that the apical ECM protein, dumpy, regulates the shape of the wings and legs through its distribution pattern (7). Dumpy has been shown to connect epithelial sheets to the cuticle (7, 8). In the wing, the long-range force pattern is regulated by the contraction of neighboring tissues and the expression pattern of dumpy, leading to the elongated wing shape. Dumpy also contributes to the differences between non-lobed and lobed posterior lobes in *Drosophila* genitalia by linking cell sheets to the apical ECM (aECM) network (9). These studies suggest that regulating long-range force patterns through cell sheet connections to ECMs (including cuticles) may play a crucial role in cell sheet deformation. However, these studies have focused on relatively simple shapes, such as wings, legs, and posterior lobes, and the mechanisms regulating more complex morphologies have yet to be fully explored.

One example of a complex morphology that undergoes a significant change in appearance before and after molting is beetle horns. The head horn of the Japanese rhinoceros beetle *Trypoxylus dichotomus* is four-branched at the tip and is one of the most elaborate horn shapes. Like other beetles, it is not present in the larval surface and is formed by three steps: folded structure formation in larvae, visible protrusion formation during pupation, and remodeling in pupae (10).

Regarding pupal horn formation, previous studies have clarified how horns suddenly appear during this process. The cell sheet that forms a pupal horn is located beneath the dome-shaped larval head capsule and, at the prepupal stage, detaches from the head capsule, undergoing cell division to form a complex folded horn primordium (11–15). The horn primordium then transforms into a pupal horn by hemolymph pressure during pupation. As previously stated, the pupal horn shape primarily depends on unfolding primordial furrows (11). The relationship between the primordial furrow patterns and unfolded 3D shapes has been revealed using computer simulations (16).

In terms of horn remodeling, previous studies have primarily focused on the biological aspects of this process (13, 17–22). Most of them have studied thoracic horn remodeling in *Onthophagus* species (17–22) as some of them lose their thoracic horns during remodeling. In *T. dichotomus*, Morita 2019 has reported that different genetic mechanisms underlie the head and thoracic horn remodeling through the detailed observation of *doublesex* RNAi mutants (13). However, the remodeling process involves both biological and physical factors, and there remains a large gap between these biological mechanisms and the resulting transformation of the epithelial sheet.

The pupal and adult horns of *T. dichotomus* are roughly similar in shape; however, there are two prominent differences. First, the distal tips of the pupal horns are rounded, while those of the adult horns are sharp. Second, the stalk is polyhedral in adults and cylindrical in pupae (Figure 1 a, b). The characteristics of adult horns resemble those of tensile membrane structures, such as outdoor tarps and tents, suggesting that tension patterns are involved in epithelial deformation during remodeling, like in the formation of wings in *Drosophila*.

**Figure 1:**
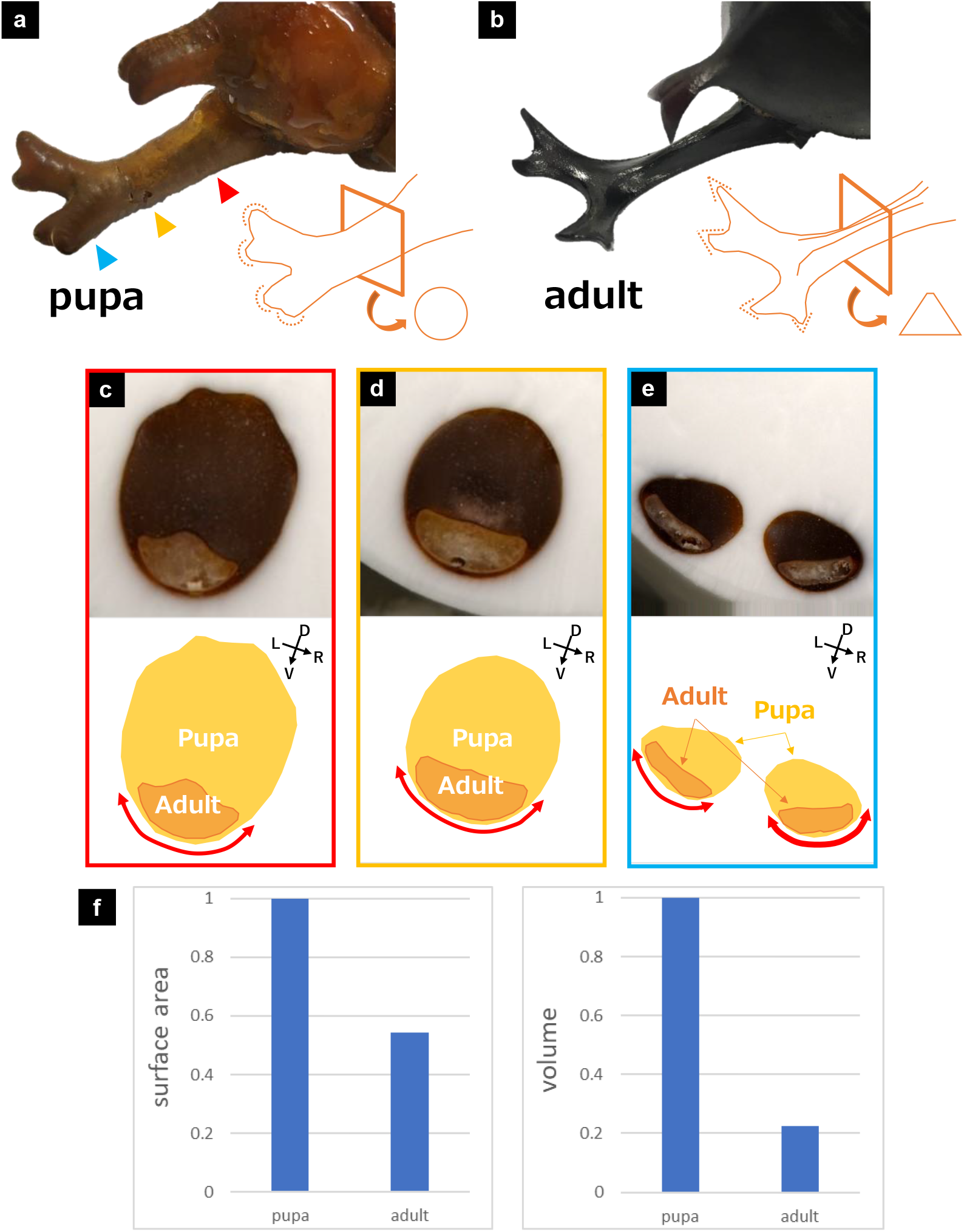
The epithelial sheet forming the adult horn decreased the surface area and the inside volume, while localized to the ventral side, during the adult horn formation. (**a,b**) Observation of a pupal horn and an adult horn. The distal tips of the pupal horn were rounded (a), while those of the adult horn were sharp (b). Also, the stalk of the pupal horn was cylindrical (a), while that of the adult horn was polyhedral (b). (**c-e**) Observation of a pupa and the adult-shaped epithelial sheet with the CoMBI method. Section images of proximal (c), middle (d), and distal (e) parts of the horn. The window colors (red, yellow, and blue) correspond to the colors of the arrowheads in Figure 1a, indicating the estimated sectioning position. The upper panels show the raw data, and the lower panels show the corresponding schematic images (yellow area: pupa, orange area: adult, bidirectional red arrow: the estimated area of the epithelia-cuticle adhesion). The epithelial sheet forming the adult horn was localized to the ventral side of the pupal cuticle over the whole horn. (**f**) Comparison of the surface area (left panel) and the inside volume (right panel) between the pupal and adult horns (n=3). The surface area and the inside volume decreased (area: ~0.5, volume: ~0.2) during the adult horn formation.

Therefore, this study focused on adhesion and contraction in the horn remodeling of *T. dichotomus*, during which a cylindrical pupal horn transforms into a polyhedral adult horn, and developed a physical simulation to reproduce this transformation. First, by detailed observation of the formation process and RNAi analysis of *dumpy* in pupae, we identified the shrinkage rate of each part of the horn, volume change, and the area of epithelia-cuticle adhesion. Next, we developed a physical simulation based on these data and demonstrated that simple physical simulations could reproduce the pupal horn remodeling. As the physical simulation was also able to reproduce the mandibular remodeling of a stag beetle to some extent, it may serve as a general model for the pupal remodeling of beetles.

## Results

### Positional and morphological relationship between a pupal horn and the adult horn

An adult beetle horn is formed inside the pupal horn and is almost complete at eclosion. Thus, we initially observed the position and shape of the epithelial sheet in a pupa using the CoMBI method (23). Figures 1c, d, and e show sectional images of the horn’s proximal, middle, and distal parts, respectively (the window colors correspond to the colors of the arrowheads in Figure 1a, indicating the estimated sectioning position), with the upper and lower panels displaying the sectional block-face and schematic images, respectively. The images showed that the pupal horns were rounded in all sections, whereas the adult horns were flattened at the distal region and more polygonal with a dorsal ridge at the proximal region. The ventral side of the adult epithelial sheet was localized to the vicinity of the pupal cuticle, suggesting that the ventral region was the site of its adhesion to the pupal cuticle.

Furthermore, micro-CT 3D mesh reconstructions (Supplementary Figure 1) revealed that an adult mesh model had approximately 50% of a pupal mesh model’s surface area and about 20% of the volume (Figure 1g, n=3).

### RNAi analysis of *dumpy*, which contributes to epithelia-cuticle adhesion

Given the images that suggested ventral epithelia-cuticle adhesion (Figure 1c-e), we investigated the effect of adhesion on epithelial sheet deformation via knockdown analysis of *dumpy*, which has been reported to be involved in epithelia-cuticle adhesion in *Drosophila* (7, 8). We injected dsRNA for dumpy RNAi into pupae and observed how the horn formation process and their final shape varied depending on the presence of adhesion.

To observe the formation process in the same pupa, we observed it by transmitting light from the dorsal and left sides (Figure 2 and Supplementary Figure 2). Figure 2a shows the changes over time in the control individuals and *dumpy* mutants. During the adult horn formation, the distal tips of the *dumpy* mutants detached and gradually moved toward the proximal side (Figure 2a, lower lane, blue arrowheads), whereas those of the control individuals stayed at the distal tips of the pupal horn (Figure 2a, upper lane).

**Figure 2:**
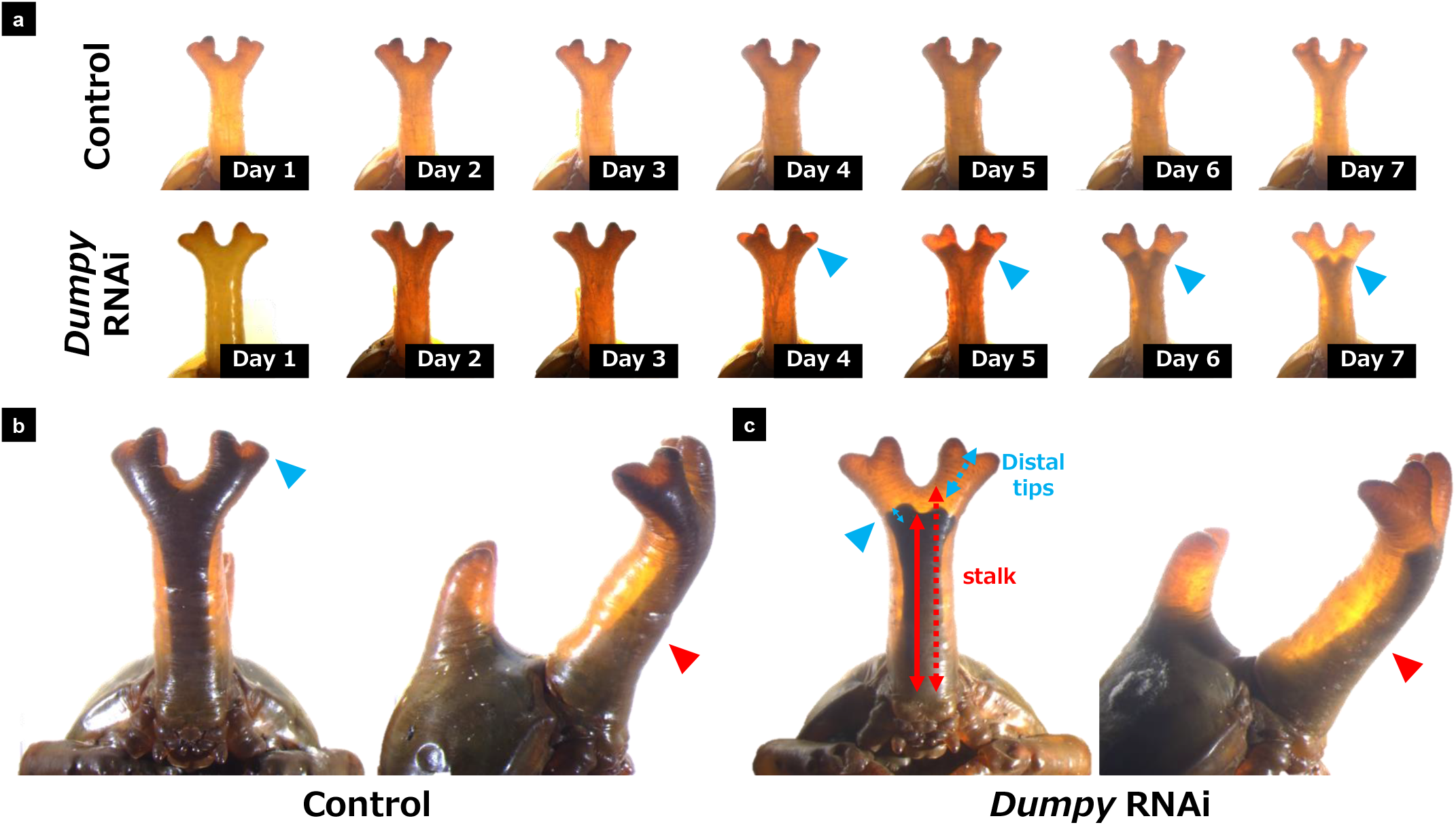
RNAi analysis of *dumpy* revealed two types of adhesion and uneven shrinkage rates between the proximal and distal parts. (**a**) Observation of the horn formation process in control individuals and *dumpy* RNAi mutants with light transmission from the dorsal side. Day X means X days after the pupation. The upper lane shows the control individual, and the lower lane shows the *dumpy* mutant. In the *dumpy* mutant, the distal tips gradually moved toward the proximal side (blue arrowheads). (**b, c**) Observation of the pupal horn that includes the adult horn (at day 20; just before eclosion) in control individuals (b) and *dumpy* RNAi mutants (c) with light transmission from the dorsal and left side. (**b**) In the control individual, the adult’s distal tips were located at the distal ends of the pupal horn (blue arrowhead), and the adult horn was stretched over the entire pupal horn. The control individual showed the localization of the adult horn to the ventral side (red arrowhead). (**c**) In the *dumpy* RNAi mutant, the adult’s distal tips were located near the distal part of the stalk (blue arrowhead). The distal tip region (blue lines, dotted: pupal, solid: adult) shrunk stronger than the stalk region (red lines, dotted: pupal, solid: adult). The *dumpy* mutant also showed the localization of the adult horn to the ventral side (red arrowhead).

Figures 2b and c show the control individual and *dumpy* mutant on day 20 when the remodeling process is completed. On day 20, the adult horn in the control individuals was stretched over the entire horn region, while the distal end in the *dumpy* mutants was located near the stalk region (Figure 2b and c, left panel, blue arrowhead). However, in both the control individuals and *dumpy* mutants, the adult horn was biased toward the ventral side of the pupa (Figure 2b and c, right panel, red arrowhead), indicating that the ventral adhesion was less susceptible to the *dumpy* knockdown. We also studied the change in the size of the stalk region relative to the distal tip region in *dumpy* mutants. The stalk region is defined as the part from the root of the horn to the bottom of the central groove, which is the first branching point (14). The distal tip region is defined as the part from the post-bifurcation constriction to the distal tip midpoint. The size comparison revealed that the distal tip region shrank more strongly than the stalk region (Figure 2b, left panel), indicating an uneven shrinkage.

### Settings of the physical simulation

The results of the biological experiments are summarized in Figure 3a. We extracted three factors involved in adult horn formation: epithelia-cuticle adhesion, uneven shrinkage, and volume reduction. The first factor can be divided into the distal *dumpy* RNAi-sensitive adhesion and the ventral *dumpy* RNAi-insensitive adhesion. Based on these factors, we developed a physical simulation to investigate whether deformation could be reproduced from the pupal mesh model. In addition to these factors, we introduced smoothing effects into the simulation, as pupae have a few wrinkles and adults do not. This is based on the idea that epithelial cells actively smoothen their surface (Figure 3b). In the simulation, the energy *U* was calculated for the mesh model, and energy minimization was performed with respect to the vertex position using the steepest descent method. The energy calculation was as follows:

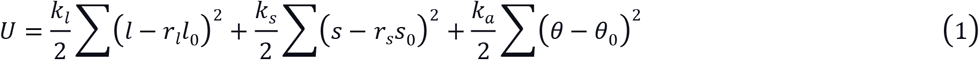

where the elastic energy was calculated for each edge length *l*, facet area *s*, and dihedral angle *θ* (*k_l_, k_s_*, and *k_a_* denote the coefficient for each elastic energy), and *l*_0_ and *s*_0_ denote the initial state of each edge length and facet area, respectively. *θ*_0_ was set as 0 radian. To express epithelial sheet shrinkage, *l*_0_ and *s*_0_ were multiplied by *r_l_* and *r_s_* (*r_l_, r_s_* < 1). Based on the energy settings, the movement of the *i*-th vertex, whose position vector was denoted by ***r**_i_* at time t, was determined by the following equation.

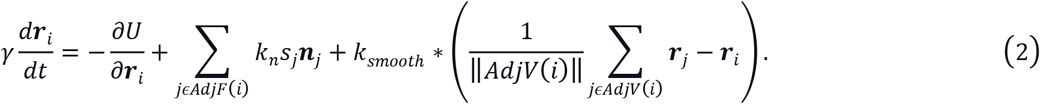

**Figure 3:**
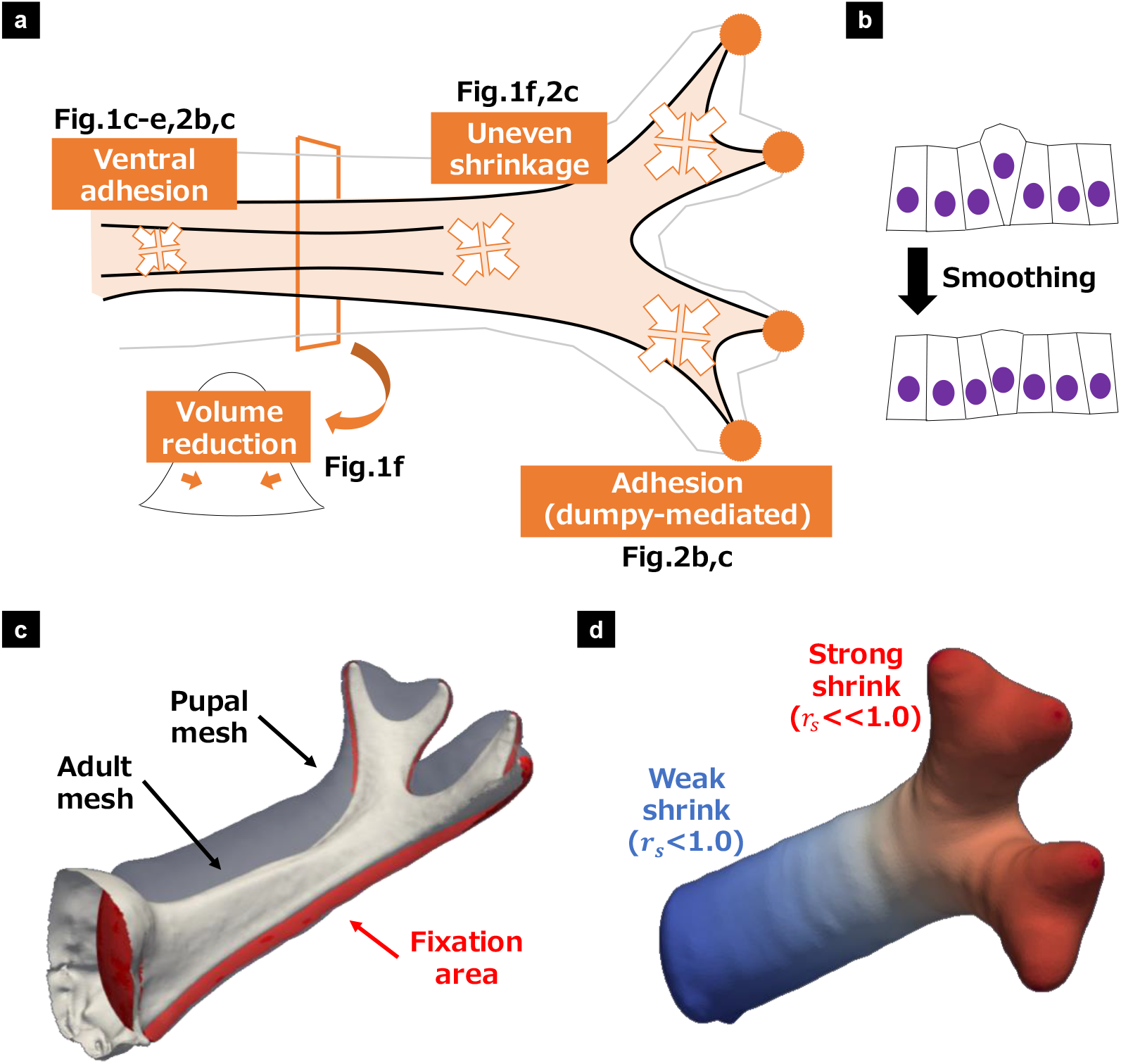
Summary of the experiments and development of the physical simulation. (**a**) Summary of the findings from the experiments. From the experimental data, we extracted three factors: ventral adhesion (light orange area) and dumpy-mediated adhesion at the distal tips (dark orange circles), uneven shrinkage of the epithelial sheet (arrows outlined in orange), and volume reduction (orange arrows). The corresponding figures are shown near each factor. (**b**) Smoothing effects of a cell sheet. Smoothing is probably due to the return of protruding cells or their elimination (e.g., by apoptosis). We implemented it as the fourth factor. The numbers in each figure show the corresponding factors. (**c**) The fixation area in the simulation. The ventral and distal tip adhesions were implemented as a simple fixation of vertices. We extracted the fixation area (red area) from the pupal mesh (transparent gray mesh) by comparing it with the adult mesh (white mesh). (**d**) The proximodistal axis in the simulation. *r_s_* (in the equation (1); the rate of facet shrinkage) in the red area was set as smaller than that in the blue area. See the supplementary information for the details of the simulation settings.

In equation (2), the first and second terms on the right-hand side express energy minimization and the negative pressure for volume reduction. The third term represents smoothing effects: it was implemented as a force on each vertex based on Laplacian smoothing, moving in the direction of the barycenter of the connecting vertices.

The distal and ventral adhesions were implemented as fixation of the vertices. The fixation area was set by distance calculation between the pupal and adult mesh models and manual selection based on the projection from the dorsal side (Figure 3c and Supplementary information 1). For uneven shrinkage, we introduced the proximodistal axis into the simulation, and *r_s_* was set smaller in the distal area than in the proximal area so that the distal area shrunk more strongly (Figure 3d and Supplementary information 2).

### The physical simulation revealed each factor’s contribution and the generality of the model

The physical simulation was performed by comprehensively modifying the parameters—*r_s_* (shrinkage ratio of the facet area), *k_a_*(stiffness of the surface), and *k_n_* (intensity of the negative pressure) — which successfully resulted in reproducing the horn remodeling (Figure 4a and Supplementary Figure 3). In particular, the simulated and actual adult horns were similar in that they were flattened in the distal region and had a ridge structure with a slope on the dorsal side of the stalk (Figure 4b).

**Figure 4:**
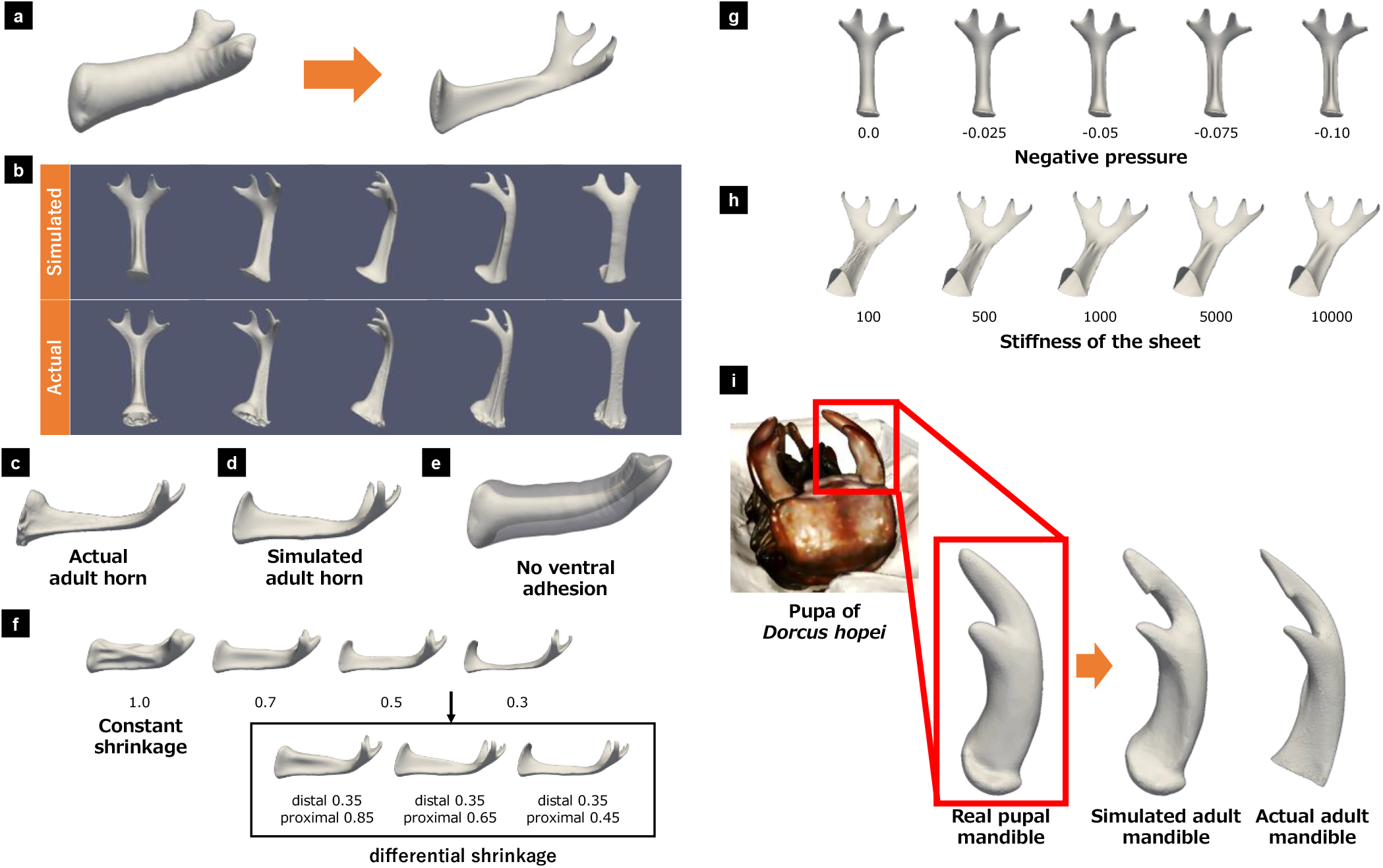
The simulation revealed each factor’s contribution to the shape determination and the generality of the simulation. (**a**) The input (left) and output (right) of the simulation. The simulation transformed the pupal mesh model into an adult-like horn. (**b**) The simulated and actual adult horns were compared from various angles (upper lane: simulated adult horn, lower lane: actual adult horn). From all angles, their shapes were similar. (**c-h**) Shape changes when the parameters and settings were varied. (**c, d**) The actual (c) and simulated (d) adult horn viewed from the right side. (**e**) Without the ventral fixation (only the distal tips were fixed), the simulated horn became a thin cylindrical shape. (**f**) The simulated horns with various shrinkage rates. The upper lane shows the simulated horns with uniform shrinkage rates (*r_s_*: position-independent). The value of *r_s_* is shown under each horn. The lower lane shows the simulated horns with uneven shrinkage rates (*r_s_*: proximodistal position-dependent). The values under each horn are the values of *r_s_* in the most distal and proximal areas, respectively. With uneven shrinkage rates, the sloped dorsal ridge appeared (lower lane). (**g**) The intensities of negative pressure were varied. The values of *k_n_* in the equation (2) are shown under each horn. In the absence of negative pressure (leftmost), the dorsal shape was arched. As the negative pressure increased (left to right), the ridge structure appeared and gradually became sharper. (**h**) The shape changes with various stiffness of the sheet. The values of *k_a_* in the equation (1) are shown under each horn. When the sheet was soft (leftmost), small wrinkles existed on the surface, although the rough structures were the same. The wrinkles disappeared as the sheet became harder (left to right). (**i**) Validation of the simulation with the mandible formation of the stag beetle *D. hopei*. The leftmost image shows the 3D data of a stag beetle. The pupal mandible mesh model was extracted from the mesh (surrounded by red lines). The simulated result was compared with the actual adult mandible (rightmost, the data was acquired with the different individuals).

To investigate the role of each factor in remodeling, we varied the factors to observe what shape changes appeared (Figure 4 c-h). Without the ventral adhesion (only the distal tips were fixed), the final shape was an elongated cylindrical shape without the typical adult polyhedral structure (Figure 4e). Without shrinkage, the final shape exhibited large folds and lacked the polyhedral structure (Figure 4f, upper lane, leftmost). When the entire mesh shrunk uniformly, the resultant shape gradually flattened, particularly in the distal region, with stronger shrinkage (Figure 4f, upper lane, left to right). However, uniform shrinkage did not generate the sloped dorsal ridge structure in the stalk region. Uneven shrinkage along the proximodistal axis caused the sloped dorsal ridge structure (Figure 4f, lower lane). With the fixed distal shrinkage rate, the size and slope of the dorsal ridge structure became slimmer and gentler as the proximal part shrunk stronger (Figure 4f, lower lane, left to right). The intensity of negative pressure was also varied. In the absence of negative pressure, an arch structure instead of a ridge structure appeared in the stalk region (Figure 4g, left), and as the negative pressure increased, a ridge structure appeared on the dorsal surface, which gradually became sharper (Figure 4g, left to right). The last factor that was varied was the stiffness of the sheet. When the sheet was soft, small folds appeared, especially in the stalk region, although the shape of the ridge was roughly similar to that of the actual adult. As the sheet became stiffer, the folds disappeared, and only the dorsal ridge structure remained (Figure 4h, left to right).

Finally, the mandible formation of the stag beetle *Dorcus hopei* was reproduced using the simulation (Figure 4i) to determine its applicability to other beetles. The 3D data of the pupa were acquired with a 3D scanner (Figure 4i, leftmost image), and the pupal mandibular mesh model was extracted from it. After the proximodistal axis and fixation area were set manually (Supplementary Figure 4), the physical simulation was applied to the mesh model, which successfully reproduced the adult form (Figure 4i, center and rightmost image).

## Discussion

In this study, we investigated the mechanisms of horn remodeling in *T. dichotomus* using physical simulations, which were applied to both the horns of *T. dichotomus* and mandibles of *D. hopei*. Many insects possess polyhedral parts; for instance, many other beetles have polyhedral-like horns. In general, insects have polyhedral-like claws (24). Because morphogenesis by “adhesion and shrinkage (in the volume and surface)” can generate polyhedral shapes, it might underlie their morphogenesis, and our simulation would help investigate them.

The simulations produced sharp protrusions when only the distal parts were fixed and the intensity of negative pressure was low relative to the surface shrinkage (Supplementary Figure 5). The polyhedral shapes and sharp protrusions generated by the “adhesion and shrinkage” mechanism are complementary to the rounded structures caused by the “furrow formation and unfolding” mechanism, indicating that both are needed to explain various exoskeletal shapes.

Previous studies have investigated the remodeling process from a biological perspective (13, 17–19). However, studying the remodeling process solely from a biological perspective is challenging, as it involves both biological and physical factors. In this study, we focused on epithelial sheet deformation and organized the contributing factors to provide insight for future biological research. Among the factors, “epithelia-cuticle adhesion” and “uneven shrinkage” could be studied biologically.

### Adhesion mechanisms

Our RNAi data suggest dumpy-dependent adhesion at the distal tips, comparable with dumpy expression in the *Drosophila* leg and antennae formations (7). In contrast, *dumpy* RNAi-insensitive ventral adhesion suggests other mechanisms. As well as the difference in the expression levels of dumpy, other ECM proteins (e.g., Obst) may explain this phenotype. The spatiotemporal regulation of apolysis may also be contributory because the type and level of chitinase expression differ depending on the insect’s stage and tissue (25).

### Shrinkage mechanisms

Like the complete resorption of a pupal horn in *Onthophagus binodis* and *O. taurus* (18), programmed cell death (PCD) might cause epithelial shrinkage in *T. dichotomus*. Although PCD can partly explain the shrinkage in *T. dichotomus*, it is different from that in *O. binodis* and *O. taurus* in that the shrinkage is not a complete resorption. Some data from this study suggested mechanical regulation of patterning of the resorption. When we created new adhesion patterns by magnetic force on the pupa, a protrusion surrounded by gentle ridges appeared (Supplementary Figure 6), suggesting that mechanisms other than the proximodistal axis also affect the shrinkage.

Our series of studies have implications for genetic research on beetle morphogenesis. While previous studies have identified many genes involved in beetle horn formation (10, 12–15, 17, 19, 20, 22, 26–29), most have examined their effects on the final morphology. Based on the results of this study and our previous study (11), beetle horn morphogenesis derives primarily from the “furrow formation” and “adhesion and shrinkage” processes. Therefore, further investigation into when and how these genes influence morphogenesis is important for improving our understanding of beetle morphogenesis.

Comparing the morphogenetic mechanisms of the head horn of *T. dichotomus* with those of the *Drosophila* leg and wing, the formation and unfolding of the furrows, followed by the generation of tension by adhesion, are common features. However, the underlying biological mechanisms differ in some respects. Unfolding furrows is mainly a physical process (internal hemolymph pressure) in beetles (11), whereas it is primarily a biological process (cell shape change and rearrangement) in Drosophila (30–34). The source of tension is thought to be the contraction of neighboring tissues in *Drosophila* (7), whereas, in beetles, the contraction of the sheet is considered important.

Regarding adhesion, both *Drosophila* and *T. dichotomus* use the same anchoring protein dumpy, but different mechanisms may mediate the ventral adhesion in the head horn remodeling. Despite the differences in biological mechanisms, since the mechanical mechanisms of “unfolding furrows” and “tension generation by adhesion” are common, they may be universally applicable to exoskeletal animals.

In the morphogenesis by “adhesion and shrinkage,” 2D adhesion patterns in the epithelial sheet control the 3D shape after shrinkage. This mechanism is similar to morphogenesis by “unfolding furrows,” in which 2D furrow patterns control the 3D shape after unfolding. Thus, both mechanisms could link 3D morphogenesis with the known genetic patterning mechanisms.

## Materials and Methods

### Insects

The beetle larvae were purchased and kept according to our previous studies. Briefly, commercially purchased last instar (third instar) larvae of the Asian rhinoceros beetle *Trypoxylus dichotomus* were kept individually in 1 L or 800 mL plastic bottles filled with rotting wood flakes at 10–15 °C to suspend their development. Larvae were moved to 25 °C to restart their development before the experiments and/or observations. Bottles were checked daily in order to record the date of pupation. Pupae were weighed before each experiment.

Larvae of stag beetle *Dorcus hopei* were purchased from Tsukiyono-kinoko-en (Gunma, Japan) and were kept in 800 mL Kinshi bottle (Tsukiyono-kinoko-en, Gunma, Japan) until pupation. The male pupa was fixed with FAA for three hours and replaced with 70% ethanol.

### Acquisition of the 3D data with micro-CT

3D information of a pupa that was just before eclosion was obtained by micro-CT, because the epithelial sheet in the pupa had cuticles on the surface, causing a clear contrast in CT values and less susceptible to deform by freeze-drying.

The frozen sample of pupa was freeze-dried. Then, dried samples were scanned using a micro-CT scanner (Skyscan1172, Bruker, USA) following the manufacturer’s instructions. The serial images were processed with ImageJ (Fiji)(35). From the serial images, meshes were reconstructed with 3D slicer software(36).

### Acquisition of the 3D data of a pupal stag beetle with a 3D scanner

The 3D data of the fixed sample was acquired with a 3D scanner (SOL 3D scanner, Global Scanning, Denmark). The mesh was reconstructed with the SOL software.

### Observation of a pupa with CoMBI

Since the epithelial sheet before secretion of the cuticles is susceptible to artifacts from drying and fixation, we employed the CoMBI method, in which biological samples were directly frozen and observed with sectioning, to obtain the 3D information of a pupa several days after pupation.

Pupae were anesthetized on ice before dissection. The dissected pupal head was mounted in OCT compound (Sakura Finetek, Tokyo, Japan). The frozen block was sectioned with a cryostat (Leica CM1850, Leica Microsystems K.K., Tokyo, Japan), and pictures of the block surface were automatically taken by a digital camera (Nikon D810, Tokyo, Japan) after every section (20 μm). The chamber temperature was set at −15 °C.

### Gene knockdown via RNAi

We searched for the ortholog mRNA sequence from the RNAseq database of T. dichotomus (PRJDB6456) using D. melanogaster sequences as a query via the tblastn program(13). Amplification of target gene sequence for making template, synthesis of dsRNA and injection for RNAi were performed as described in our previous study. 5 μg of dsRNA of each target gene was injected into pupae. The primer sequences for amplifying target gene sequence: F-ACCTGTCGACCAGAACCAAC, R-GCAGGAACAAGAAGCCTGTC.

### Mesh processing

The acquired mesh was cleaned with Meshlab(37). After cleaned, the mesh was processed with the Poisson remeshing filter with Meshlab, to generate a single-layered mesh with no holes or tunnels. Then, meshes were processed with the Quadric Edge Collapse Decimation filter, to reduce the number of vertices and facets. The information of each mesh is shown in Supplementary Table 1.

### Physical simulation

To simulate the development of an adult horn, energy minimization was performed with respect to the vertex position using the steepest descent method, as described in our previous papers. The numerical calculation was performed with sundials(38). See the Result section for the details of the energy settings and the force calculation. The details of the settings of the proximodistal axis and the fixation area are written in the supplementary information. The simulated data was visualized with ParaView(39). The parameters for each simulation are shown in Supplementary Table 2.

## Acknowledgement

We appreciate Dr. S. Morita for his advice in the beads injection into pupal horns. This work was supported in part by JSPS KAKENHI Grant Number 15H05864 (to HG, YI, and SK), 20A306 (to HG, YI, and SK), 22J10710 (to KM). KM was supported by Grand-in-Aid for JSPS Fellows (DC2) and the ANRI Fellowship. HA was also supported by Grand-in-Aid for JSPS Fellows (DC2).

## Author Contributions

Conceptualization: K.M. and S.K. Formal analysis: K.M. Funding acquisition: K.M., H.G., Y.I., and S.K. Methodology: K.M. Resources: H.G. and H.A. Software: K.M. and Y.I. Supervision: S.K. Writing—original draft: K.M. and S.K. Writing—review and editing: K.M., H.A., H.G., Y.I., and S.K.

## Data Availability

All study data are included in the article and/or Supplementary files. The codes used in this paper will be made available on the Bitbucket after acceptance.

## Supplementary Materials

Supplementary Figure 1-6

Supplementary Table 1, 2

Supplementary Information 1, 2

