## Supplementary Materials for "Mechanical models affecting beetle horn remodeling"

Supplemenatry Figure 1: Reconstruction of the pupal and corresponding adult meshes.

Supplemenatry Figure 2: Time-lapse images of *Dumpy* RNAi and Control (viewed from the right side).

Supplemenatry Figure 3: Comparison between the actual and simulated adult horn meshes.

Supplemenatry Figure 4: The simulation settings for the mandible formation of a stag beetle.

Supplementary Figure 5: Epithelial sheet deformation in the beetle horn morphogenesis.

Supplementary Figure 6: Administration of magnetic beads to induce novel adhesion patterns.

Supplementary Table 1: Information of the meshes for the physical simulation.

Supplementary Table 2: The parameters for the physical simulation.

Supplemenatry Information 1: Determination of the fixation area for the beetle horn formation.

Supplementary Information 2: The way to set the proximodistal axis for the physical simulation

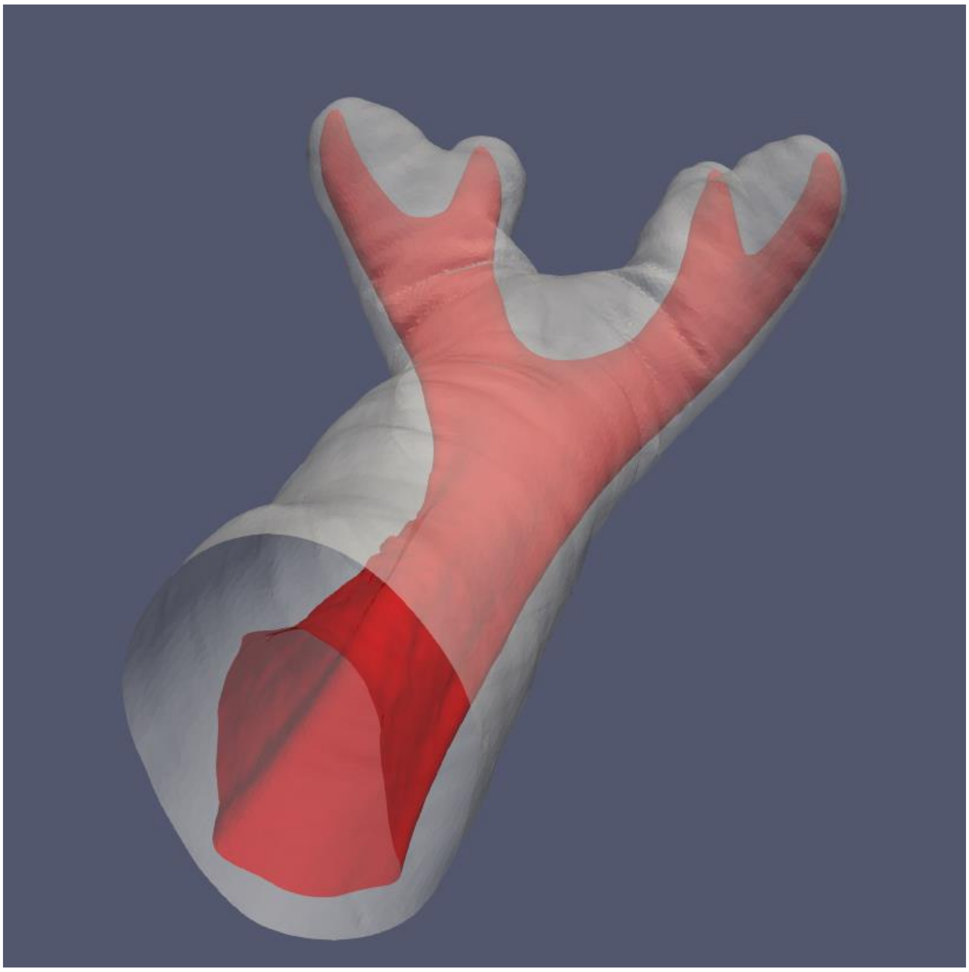

**Supplementary Figure 1: Reconstruction of the pupal and corresponding adult meshes.**

The pupal (white transparent mesh) and corresponding adult mesh (red transparent mesh) were reconstructed from the data of microCT. These meshes were used to calculate the surface area and the inside volume and to perform physical simulations.

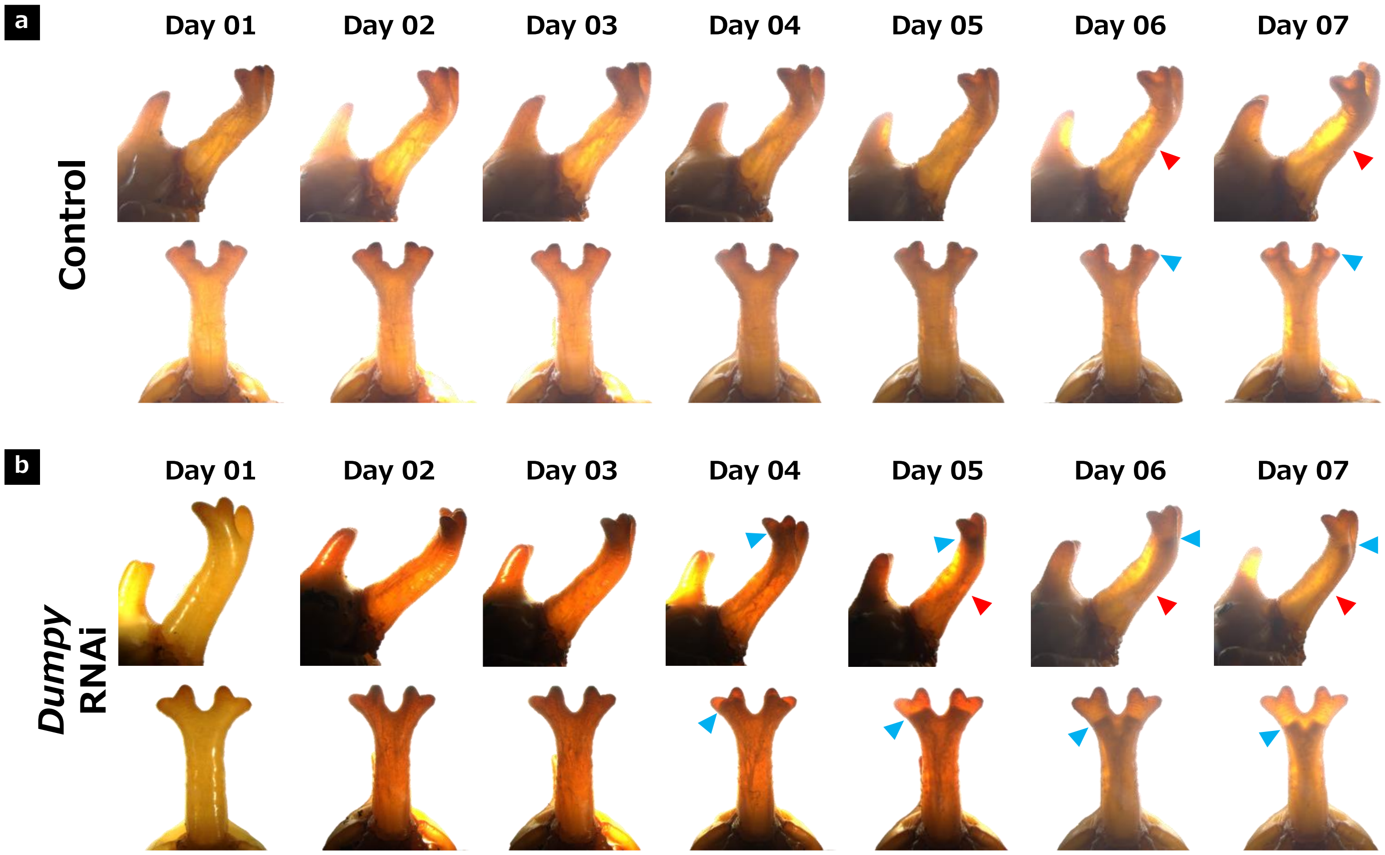


**Supplementary Figure 2: Time-lapse images of *Dumpy* RNAi mutants and control individuals (viewed from the right side).**

We performed an RNAi analysis of *Dumpy*, and observed the horn formation process in control individuals and *Dumpy* RNAi mutants with light transmission from the left and dorsal sides.

(a) The horn formation process of the control individual. The upper lane shows the images viewed from the right side, and the lower lane shows the images viewed from the ventral side.

(b) The horn formation process of the *Dumpy* RNAi mutant. The upper lane shows the images viewed from the right side, and the lower lane shows the images viewed from the ventral side.

The localization of the adult horn to the ventral side was maintained over the entire formation process both in the control individuals and the *Dumpy* mutants (red arrowheads). Blue arrowheads indicate the estimated distal tips.

Figure 2a shows the changes over time in control individuals and *Dumpy* mutants. During the adult horn formation, the RNAi mutants' distal tips were detached and gradually moved toward the proximal side (Figure 2a, lower lane, blue arrowheads). In contrast, the distal tips of control individuals extended to the entire pupal horn (Figure 2a, upper lane).

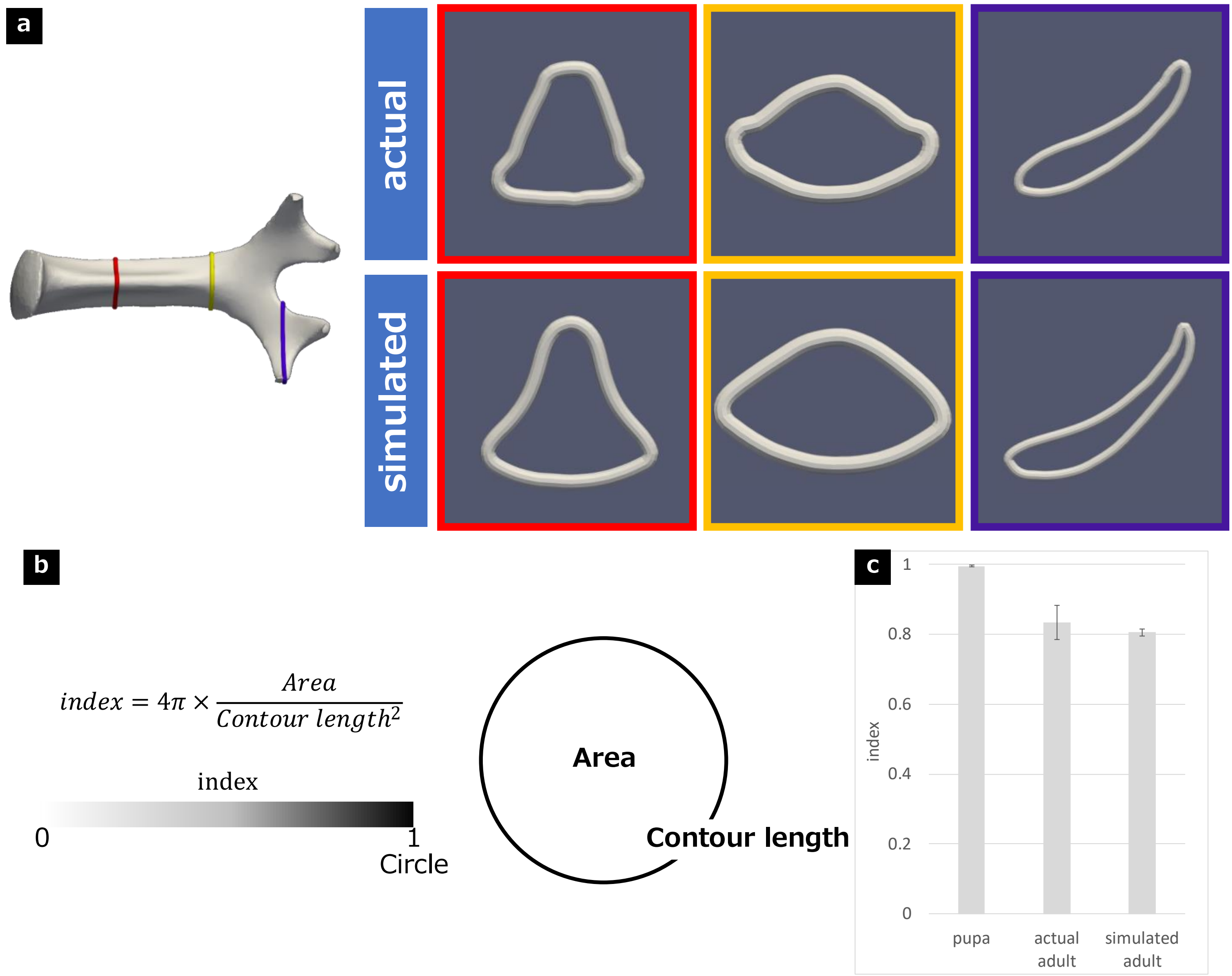


**Supplementary Figure 3: Comparison between the actual and simulated adult horn meshes.**

(a) Section images of the actual and simulated adult horn. The red, yellow, and blue windows, respectively, show the proximal, middle, and distal section images. The cross-sectional comparison showed similarities in the ridge structure and the appearance of lateral extrusions in the proximal part. In the distal part, the simulated horn was similar to the actual horn in that both had a flattened structure with a concave dorsal surface, and in the middle part, they had a similar rhomboidal shape.

(b) The way to quantify the shape of the stalk. The index was calculated with each section by the upper left equation. The index takes a value between 0 and 1. The closer the value is to 1, the closer the shape of the section is to a circle. Since the section of the pupal stalk was similar to a circle, the index value was close to 1.

(c) Quantified values were compared for pupal, actual adult, and simulated adult horns. The simulated adult horns were close to the actual adult horns. The quantification was performed with three samples, respectively. The average value of the index over the stalk was calculated with each sample. The graph shows the mean and the standard deviation.

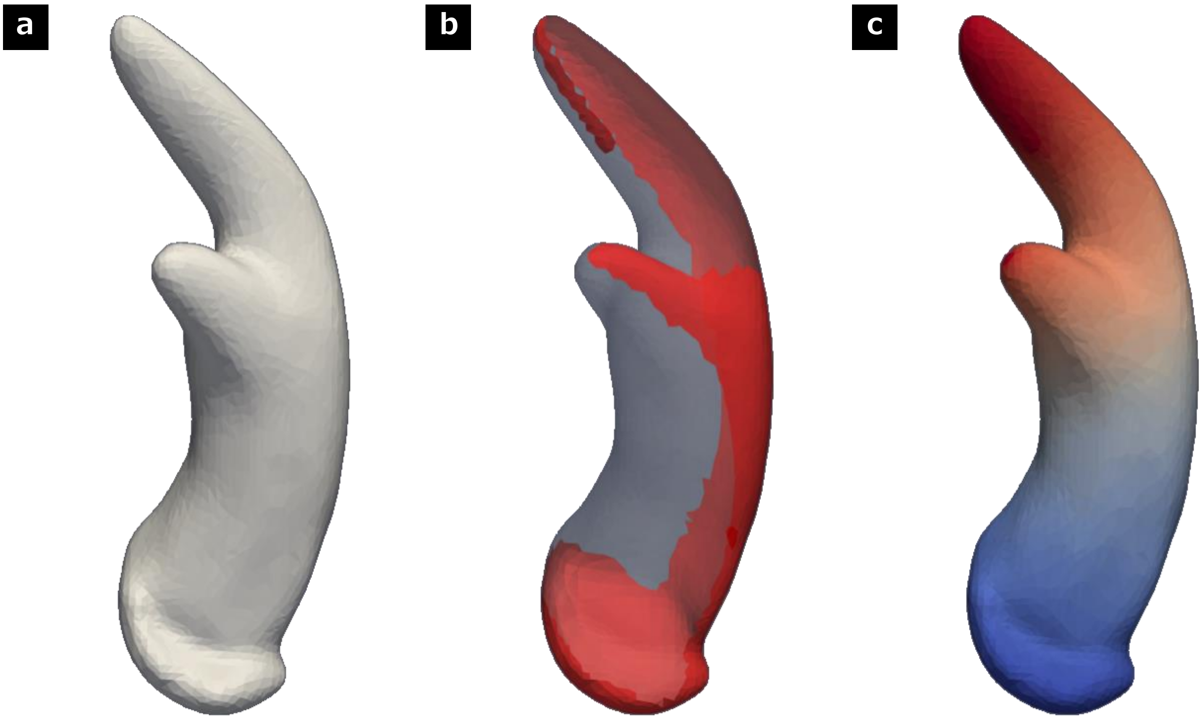


**Supplementary Figure 4: The simulation settings for the mandible formation of a stag beetle.**

(a) The pupal mandible mesh of a stag beetle *Dorcus hopei binodulosus*.

(b) The fixation area in the simulation. The gray transparent mesh shows the pupal mandible mesh, and the red area shows the fixation area. The fixation area in this simulation was determined entirely manually, base d on observing the adult shape.

(c) The proximodistal axis in the simulation. The red area was treated as the distal area, and the blue area wa s treated as the proximal area.

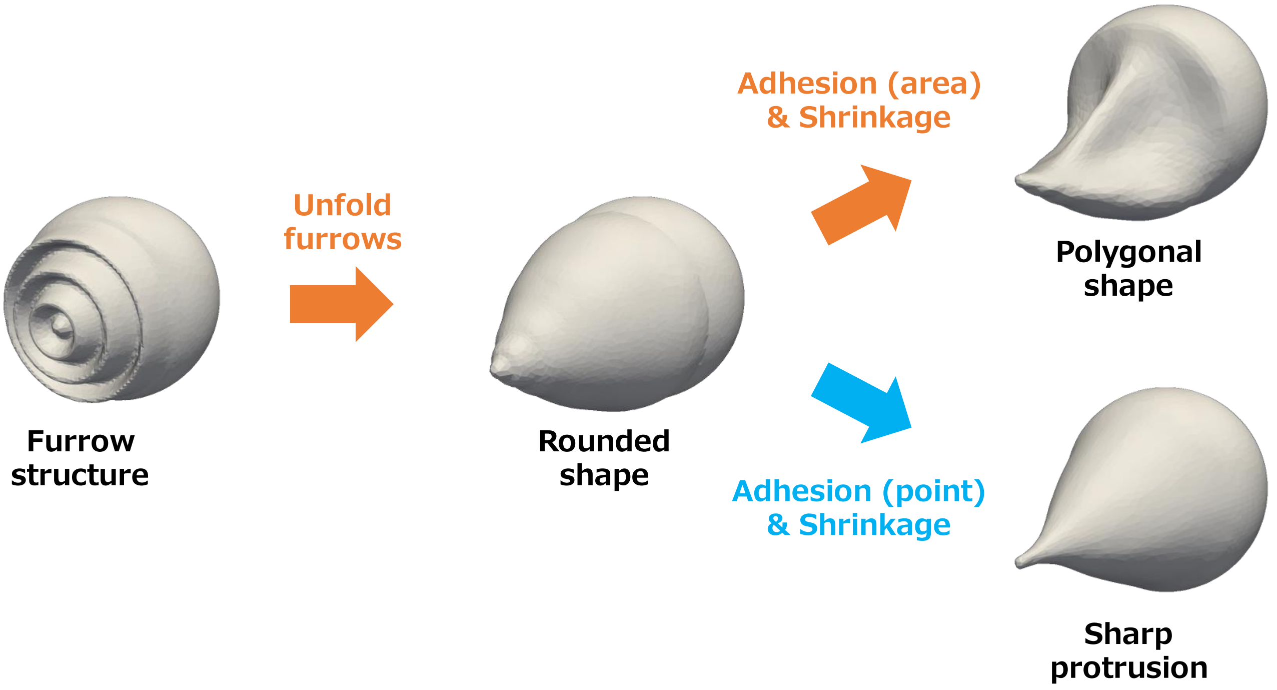


**Supplementary Figure 5: Epithelial sheet deformation in the beetle horn morphogenesis.**

The furrow structure, such as a horn primordium, is unfolded during pupation to form a pupal rounded sha pe. The rounded shape is transformed into an adult polyhedral shape by adhesion and shrinkage (orange ar rows). Using the same method, protrusions can be generated when only the distal part is fixed and the inten sity of negative pressure was low relative to the surface shrinkage (blue arrow).

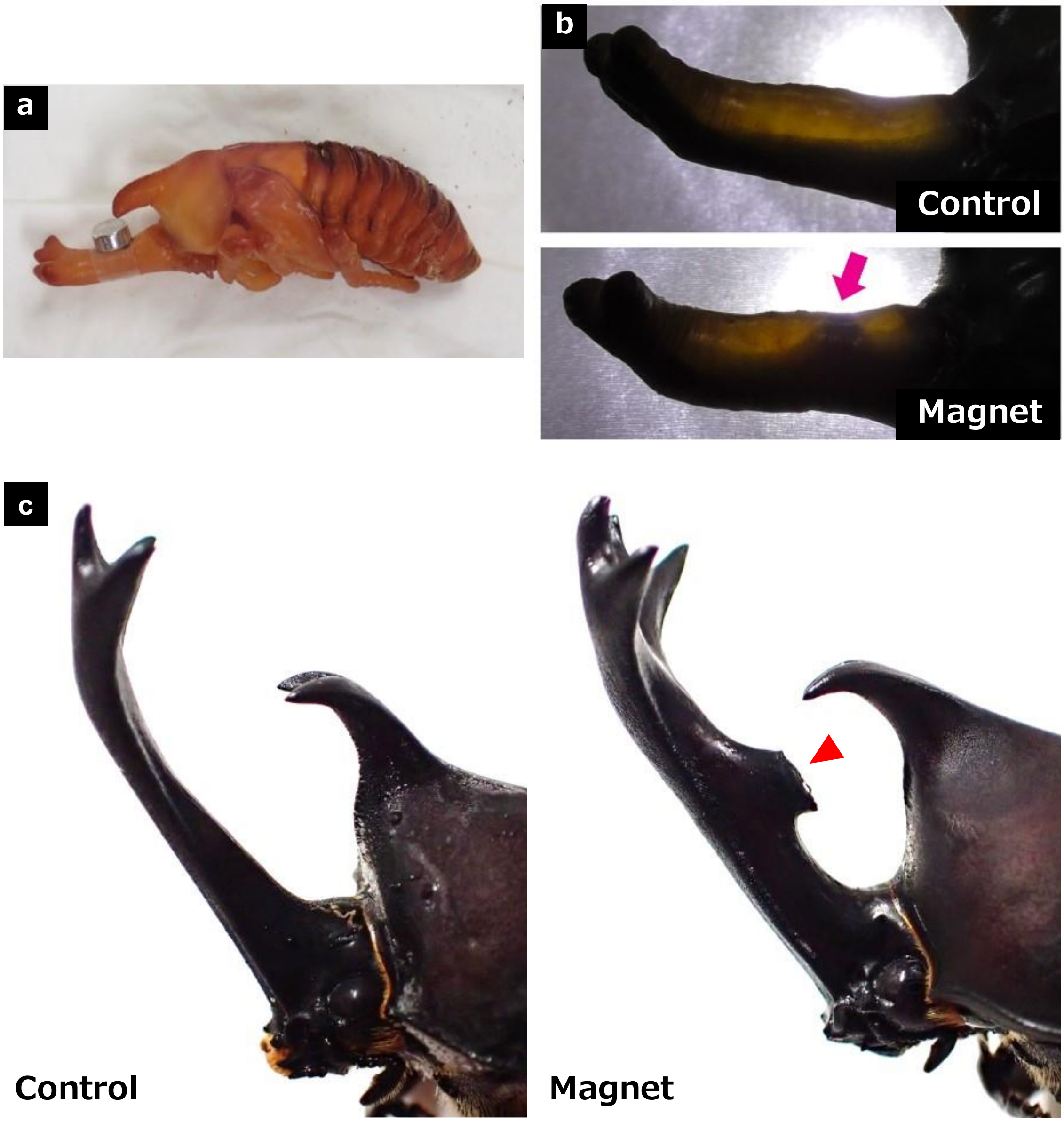


**Supplementary Figure 6: Administration of magnetic beads to induce novel adhesion patterns.**

(a) 5 μL of magnetic beads (SupraBead-Silica 4-6 μm, Recenttec, Tokyo, Japan) was injected into pupal horn by Hamilton syringe 1701N (Hamilton, Reno, NV) within 24 hours after pupation. Then, they grew while a m agnet was attached to the outside of the horn.

(b) The appearance of the deformed cell sheet in the control pupa and the magnetic beads-treated pupa wa s observed with the light transmission; new protrusions appeared in the magnetic beads-treated pupa (pink arrow).

(c) Changes in the 3D shape caused by artificial novel adhesion patterns. A new protrusive structure appear ed with the administration of magnetic beads to the pupa (right panel, red arrowhead).

**Supplementary Table 1: Information of the meshes for the physical simulation.**

| | beetle 1 | beetle 2 | beetle 3 | stag beetle |
|---|---|---|---|---|
| Number of vertices | 16522 | 12288 | 4693 | 2619 |
| Number of facets | 33040 | 24572 | 9382 | 5234 |

**Supplementary Table 2: The parameters for the physical simulation.**

The parameters in each simulation are shown in the following table. In the table, $T_{max}$ is the finish time of th e simulation.

| parameter | beetle 1 | beetle 2 | beetle 3 | stag beetle |
|---|---|---|---|---|
| $k_l$ | 2e+1 | 2e+1 | 2e+1 | 2e+1 |
| $k_s$ | 5e+1 | 5e+1 | 5e+1 | 5e+1 |
| $k_a$ | 1e+4 | 1e+4 | 1e+5 | 2e+4 |
| $k_n$ | -0.07 | -0.07 | -0.15 | -0.1 |
| $k_{smooth}$ | 1e+2 | 1e+0 | 1e+1 | 1e+1 |
| $T_{max}$ | 2.00 | 2.00 | 0.25 | 0.30 |
| $r_{s_{distal}}$ | 0.35 | 0.35 | 0.40 | 0.60 |
| $r_{s_{proximal}}$ | 0.75 | 0.55 | 0.80 | 1.00 |

**Supplemenatry Information 1: Determination of the fixation area for the beetle horn formation.**

A pupal mesh and the corresponding adult mesh were prepared to extract the fixation area from the pupal mesh. In this paper, the 3D meshes were reconstructed from micro-CT data.

The first step in the extraction process was to estimate the closeness between the pupa and the correspondi ng adult. To estimate the closeness, the minimum distance between the pupal and adult mesh was calculate d.

In the following description, $vp_i$ denotes the $i$-th vertex belonging to the pupal mesh, and $va_j$ denotes the $j$ -th vertex belonging to the adult mesh. To calculate the minimum distance between the pupal and adult me shes, we first calculated $d_{i,j}$, the distance between $vp_i$ and $va_j$. $d_{i,j}$ was calculated for all $j$ to search the mini mum value of $d_{i,j}$. Since the minimum value depends on $i$ (the position on the pupa), it was expressed as $minimum\_distance_i$ and considered as the closeness between the pupa and the corresponding adult.

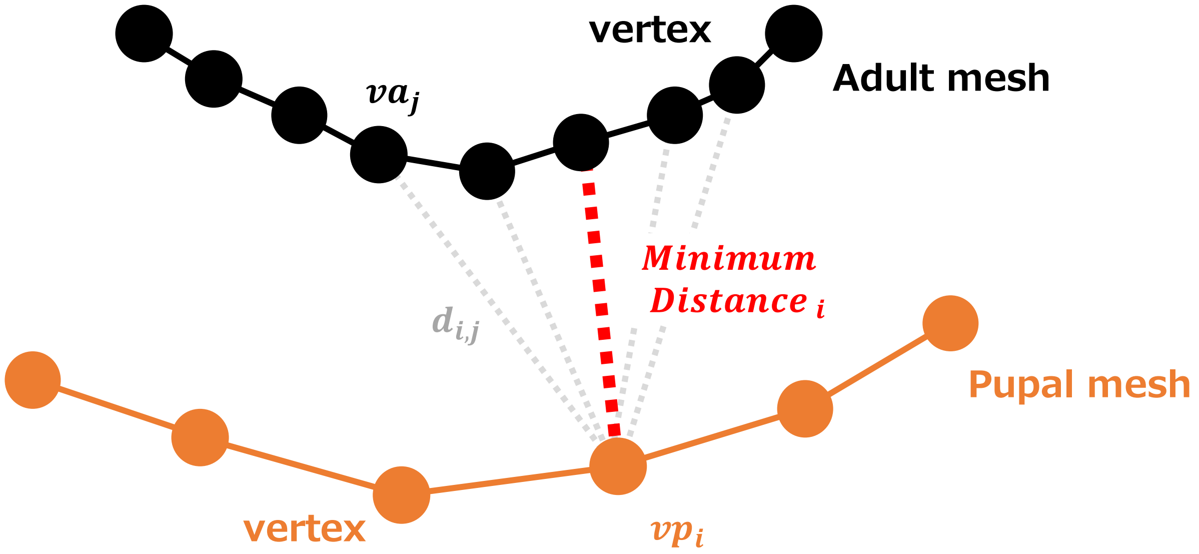


**Calculation of the minimum distance between the pupal and adult vertices (≈closeness).**

The following figure shows the heatmap of $minimum\_distance_i$. The red area indicates a small value $minimum\_distance_i$, and the blue area indicates a large value of $minimum\_distance_i$.

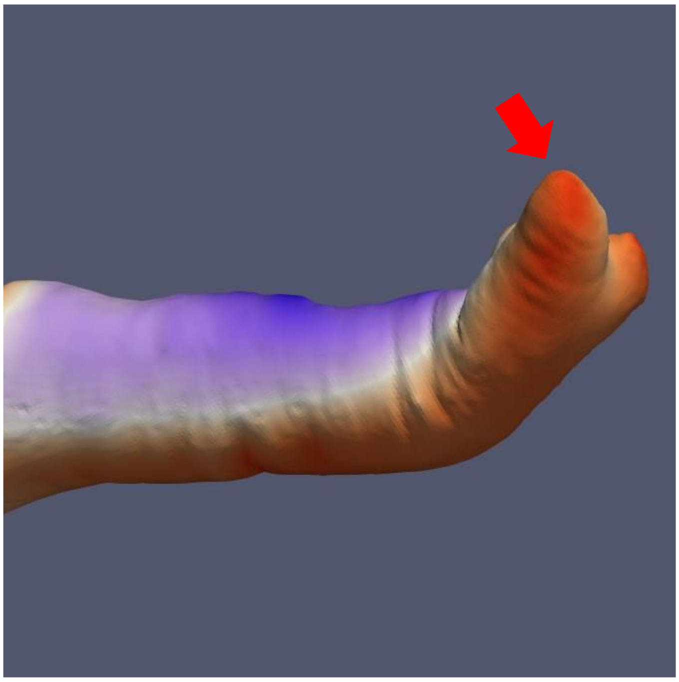

**Heatmap of $\boldsymbol{minimum\_distance_i}$.**

Then, we extracted the vertices whose $\boldsymbol{minimum\_distance_i}$ was smaller than the threshold.

The method enabled us to obtain most of the area where the adult cell sheet adheres to the pupal cuticle. H owever, the extracted mesh has an excess area that is not on the ventral side. The excess area is located in t he boundary of the area, especially in the tip area, because the shape is close to the sphere (red arrows in th e figure above).

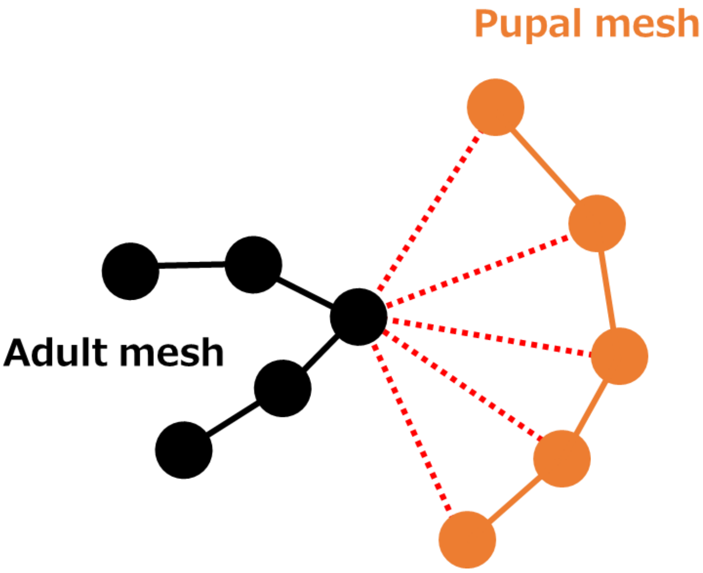


**The values of $\boldsymbol{minimum\_distance_i}$ are almost the same in the sphere-like area.**

So, we manually trimmed off the excess area from the extracted mesh to determine the fixation area.

In the trimming process, the area that protrudes from the corresponding adult horn when viewed from the dorsal side was trimmed as the excess area.

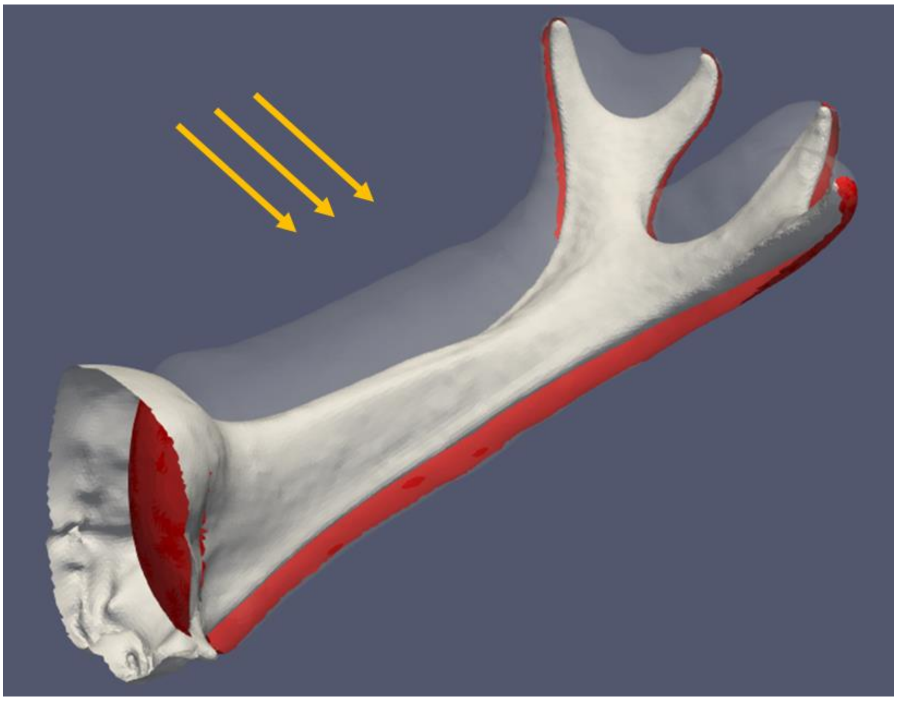

**Manual trimming of the excess area.**

**Supplementary Information 2: Generation of the proximodistal axis for the physical simulation.**

To generate the proximodistal axis, we developed a diffusion simulator.

In the simulator, each facet had information on the concentration of the virtual diffusion molecule. The dista l tips were sources of the molecule (the red circles in the following figure), and they diffused to the proximal side (the blue dotted line in the following figure).

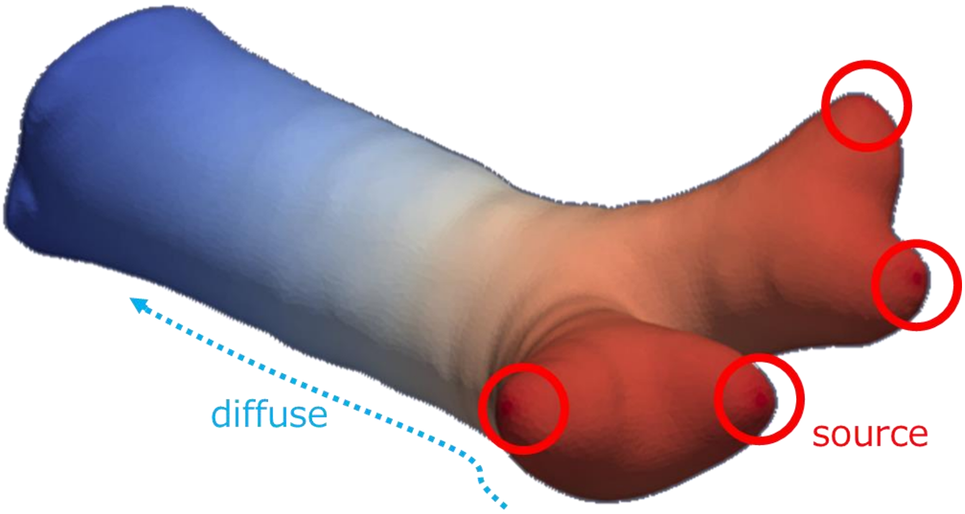


**Generation of the proximodistal axis.**

After a sufficient diffusion calculation (some time after the concentration became greater than 0 at the most proximal location), the proximodistal position of $i$-th facet $pd\ pos_i$ was calculated as

$$pd\ pos_i = \log(C_i).$$

Then, the value of $r_s$ (shown in Figure 3d) of $i$-th facet was calculated by the following equation.

$$r_{s\,i} = \frac{pd\ pos_{max} - pd\ pos_i}{pd\ pos_{max} - pd\ pos_{min}} \times \left( r_{s\,proximal} - r_{s\,distal} \right) + r_{s\,distal}$$

In the equation, $pd\ pos_{max}$ and $pd\ pos_{min}$ are the maximum and minimum values of $pd\ pos_i$, respectively. $r_{s\,distal}$ and $r_{s\,proximal}$ are the given values of $r_s$ in the most distal and proximal area.
